# Phenotyping and molecular marker analysis of WH1105 and Kharchia 65 backcrosses and F_4_ progenies for salinity tolerance

**DOI:** 10.1101/586941

**Authors:** Varsha, Shikha Yashveer, Vikram Singh, Swati Pratap

## Abstract

Soil salinity is a worldwide adverse environmental factor for crop productivity and quality in arid, semiarid and coastal areas. In India, approximately 8.5 million hectare of land area is affected by high salinity (EC ≥ 5 dS m^−1^). Development of salinity tolerant varieties through marker assisted breeding is most efficient and effective strategy for management of soil salinity. WH 1105 is widely cultivated wheat variety with many agronomically superior qualities but is affected by soil salinity. Two genes (*Nax1* and *Nax2*) for salinity tolerance were introgressed from Kharchia 65 into the genetic background of WH 1105 through marker assisted backcross breeding. BC_1_F_3_, BC_2_F_2_ and F_4_ generations of the cross WH1105 x Kharchia 65 were evaluated for various morphological traits under initial salt stress condition. On the basis of phenotypic and genotypic variations 44 high yielding plants were selected from the cross. Out of 178 SSRs tested, 30 were found polymorphic for background selection of the foreground selected plants. Cluster tree analysis of parents and all the three generations showed that all the selected plants were inclined toward recurrent parent (WH 1105) indicating higher similarity with the recurrent parent. Four plants were selected as high grain yielding and salt tolerant. These plants could be further backcrossed with the recurrent parent to develop salt tolerant wheat lines.

## INTRODUCTION

Wheat (*Triticum aestivum* L. em. Thell) was one of the first domesticated food crops and has been the basic staple food of the major civilizations of Europe, West Asia and North Africa. Today, wheat covers major land area than any other commercial crop and continues to be the most important food grain source contributing about a fifth of total calories consumed by humans. Globally among food grains the nutri-rich wheat is largest cultivated crop across countries having area around 219 mha with an annual production of 760.3 mt in 2017-18 (FAOSTAT, 2017). In India, wheat production in 2017-18 Rabi season has made another record output of 98.61 mt with an average national all time highest productivity of 33.18 q/ha in 29.72 mha area. The national average productivity of crop increased significantly by 1.18q/ha (+3.68%) which is a major reason for the increased production.

Productivity of the crops mainly depends on the genetic efficiency of the crop, environmental conditions, soil type, biotic and abiotic stresses. Among abiotic stresses, soil salinity is a major restricting factor with limiting effects mainly on seed germination, plant vigour and crop yield. Salinity is characterized by a high concentration of soluble salts in the soil. Soils are classified as saline when the ECe is 4 dS/m or more (USDA-ARS. 2008). This definition of salinity derives from the ECe that significantly reduces the yield of most crops associated with increase in Na^+^ and Cl^−^ ions causing many disorders in plants like ion toxicity, nutritional disorders, genotoxicity,waterstress,oxidativestress,alterationofmetabolicprocesses,membrane disorganization, reduction of cell division and expansion (Carillo *et al.* 2011 and Parvaiz *et al.* 2013). Jointly, these effects reduce plant growth, development and survival.

Although wheat is considered moderately salt tolerant, increasing salinity is becoming one of the most important environmental constrains in global wheat production. To meet future grain demands, it is required to develop new resource-efficient and salt tolerant wheat cultivars. Over the years, the traditional phenotyping methods have been used to screen crops agronomic and yield performance under salinity stress. However, these methods are labor-intensive, time-consuming and are considered to be one of the major limitation in utilizing genetic information for genomic analysis (Rahaman *et al.*, 2015) and furtherance of yield improvement (Furbank and Tester, 2011). The evaluation of large populations in breeding programs has necessitated the search for traits associated to salinity tolerance that are quick, easy and economical to measure and advancement.

DNA markers linked to specific genes/quantitative trait loci (QTL) have improved the efficiency of selection, in particular for those traits that are multi-genic and are greatly influenced by environmental factors (Yamaguchi and Blumwald, 2005). Microsatellites markers have been extensively used for investigation, genetic divergence, genome and QTL mapping for salt tolerance in different crops (Liu *et al.*, 2001; Munns, 2002; Ma *et al.*, 2007 and Kurup *et al.*, 2009). Two major genes for Na^+^ exclusion, termed *Nax1* and *Nax2*, were identified in Line 149 originated from a wheat relative, *Triticum monococcum* (C68-101) (Munns and James, 2003). Utilization of the previously confirmed molecular markers for *Nax1* and *Nax2* validated for salt tolerance in Kharchia 65 has been introgressed to HD 2851 (Yadav *et al.*, 2017). Location of *Nax1* gene on chromosome 2A was mapped by QTL analysis and identification as Na^+^ transporter of the HKT gene family HKT7 (HKT1; 4) was done by RFLP, AFLP and Microsatellite markers (Lindsay *et al.*, 2004). *Nax2* was mapped to the distal region on chromosome 5AL based on linkage to microsatellite markers which coincides with the locus for a putative Na^+^ transporter, HKT1; 5 (HKT8) (Byrt *et al.*, 2007).

*Nax1* removes Na^+^ from the xylem as it enters the shoot so that Na^+^ retained in the base of the leaf, impart a high leaf sheath: blade ratio, while on the other hand, *Nax2* removes Na^+^ from the root xylem (Zhang and Shi, 2013). Since *Nax* genes are not present in modern wheat (Ashraf and Foolad, 2013) molecular markers linked with *Nax1* and *Nax2 genes* can be employed as an effective screening tool to overcoming the salinity problem which limits the wheat production.

## MATERIALS AND METHODS

### Material

The experimental plant material comprised of WH 1105 x Kharchia 65 derived BC_1_F_3_, BC_2_F_2_ and F_4_ seeds. Kharchia 65 is a salt tolerant genotype, though due to rust susceptibility it is not popular among farmers but still contributes in many breeding programs for its salinity and drought tolerance. WH 1105 is a popular wheat variety of North Western Plain Zone (NWPZ) of India with high yield potential (28.64 quintals per acre) and good agronomic traits.

### Raising of crop for Phenotyping

The back cross and F_4_ generations of WH 1105 x Kharchia 65 along with parental genotypes were raised during *Rabi* 2016-17 in the net house of the Department of Molecular Biology, Biotechnology and Bioinformatics, CCS Haryana Agricultural University, Hisar. Seeds of all the progenies were sown in the trays filled with soil and vermiculite mixture. The seeds were subjected to chloride dominated salt stress (EC ~ 8 ds/m) at the time of germination and after about 15-21 days, seeds germinated in this salt stress were transferred to pots with normal soil i.e. without any salt stress. Before transferring, the original soil of the seedlings was neutralized by washing them in water and the hoagland nutrient solution was given to the plants growing in pots in the net house after every 10 days from the sowing date. The seedlings were grown to maturity to collect data on various morphological traits.

### Genotyping

Genomic DNA was isolated from young leaf tissues of the morphologically superior BC_1_F_3_, BC_2_F_2_ and F_4_ plants using CTAB method (Saghai-Maroof *et al.*, 1984). The individual plants were screened for the presence of *Nax* loci. A total of 178 SSR primers were then used to study polymorphism among parents. The plants having *Nax* loci were subjected to amplification using polymorphic markers for background selection. Plants showing the absence of *Nax* loci but morpho-physiologically superior especially in terms of yield trait were also included for further study assuming some other salt tolerance mechanism may be working in these plants.

### Data analysis

Morphological data was analyzed using INDOSTAT software. The presence of band of DNA on agarose gel was taken as one and absence of band was read as zero. The 0/1 matrix was used to calculate similarity genetic distance using “simqual” sub-program of software numerical taxonomy and multivariate analysis system (NTSYS-pc v2.02) (Rohlf, 1993). Dendrogram was constructed by using distance matrix by the unweighted pair-group method with arithmetic average (UPGMA) sub-programme of NTSYS-pc. Principle component analysis (PCA) was done using the “PCA” sub programme of NTSYS-pc.

## RESULTS

### Mean performance of parents under control and salt stress condition

Parental wheat genotypes of the cross i.e. WH 1105 and Kharchia 65 were sown in the net house under control and salt stress conditions. The elite genotype WH1105 used as female parent in cross recorded a reduction in all morphological trait values except 100 grain weight under saline conditions as compared to normal conditions (Table 1). This reduction in trait values was less in Kharchia 65. In fact, Kharchia 65 showed an increase in values for the traits like spike length, and harvest index under salt stress as compared to normal.

**Table 1:** Mean performance of parents under control and salt stress condition.

| Traits | Mean ± S.E | | Mean ± S.E | | Per cent reduction in trait values under salt stress | | LSD @<br>P = 0.05<br>(variety) | LSD @<br>P = 0.05<br>(treatment) |
| --- | --- | --- | --- | --- | --- | --- | --- | --- |
|  | (Under Control) |  | (Under salinity) |  |  |  |  |  |
|  | WH 1105 | Kharchia 65 | WH 1105 | Kharchia 65 | WH 1105 | Kharchia 65 |  |  |
| Plant height (cm) | 74.67±1.39 | 84.42±0.59 | 69.30±1.57 | 80.87±2.09 | 7.19 | 4.21 | 0.11* | 0.04* |
| Spike length (cm) | 9.95±0.24 | 9.90±0.53 | 9.88±0.77 | 10.10±0.32 | 0.67 | -2.02 | NA | NA |
| No. of tillers /plant | 1.83±0.48 | 4.60±0.40 | 1.50±0.22 | 3.50±0.22 | 18.18 | 23.91 | 0.08* | 0.46* |
| Number of spikelets/spike | 16.33±0.67 | 15.00±0.63 | 12.67±0.80 | 13.00±0.52 | 22.45 | 13.33 | 0.44* | 0.14* |
| Number of grains /spike | 53.67±1.12 | 45.80±1.69 | 37.00±1.13 | 45.67±2.39 | 31.06 | 0.29 | 0.68* | 0.12* |
| 100 Grain Weight (g) | 3.34±0.16 | 2.35±0.05 | 3.62±0.31 | 1.85±0.06 | -8.63 | 21.27 | 0.37* | 0.49* |
| Grain yield/ plant (g) | 2.55±0.29 | 1.80±0.09 | 1.74±0.08 | 1.78±0.02 | 31.85 | 1.06 | 0.32* | 0.45* |
| Biological yield/ plant (g) | 6.99±0.63 | 7.80±0.36 | 6.25±0.33 | 7.20±0.14 | 10.52 | 7.69 | 0.1* | 0.42* |
| Harvest index (%) | 36.13±1.30 | 22.40±0.32 | 27.84±0.41 | 22.69±0.31 | 22.95 | -1.27 | 0.92* | NA |

\* LSD (Least Significant Difference) makes a direct comparison between two means from two individual groups. Difference larger than the LSD gives significant result

### Mean performance for various traits in backcrosses and F_4_ populations under initial salt stress

A total of 1190 seeds (Table 2) were sown in the trays filled with soil and vermiculite mixture. Out of these 1190 seeds, 205 plants of BC_1_F_3_, 63 of BC_2_F_2_ and 70 of F_4_ survived up to maturity and were harvested separately.

**Table 2:**
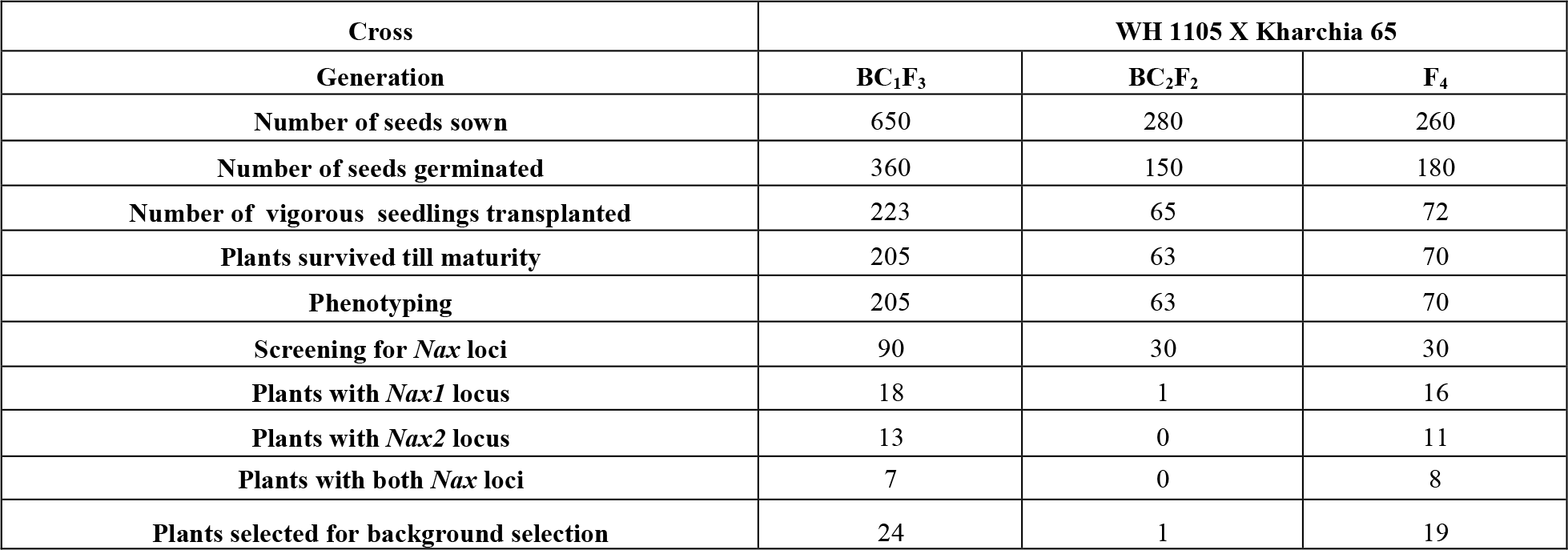
Brief details of sowing and other parameters evaluated at various stages.

### Variability parameters for various traits in backcrosses and F_4_ population

The genetic parameters were studied for the various morphological traits in different populations. A wide range of variation was observed for all the traits in parents and populations of all the generations. The PCV, GCV, heritability (H^2^) and genetic advance as 5% of mean for different traits are compiled in tables 3, 4 and 5 for BC_1_F_3_, BC_2_F_2_ and F_4_ populations, respectively. The perusal of the tables revealed that PCV was higher than respective GCV for all the traits under study. In all the three generations (BC_1_F_3_, BC_2_F_2_ and F_4_), highest phenotypic as well as genotypic coefficient of variation was recorded for grain yield per plant. Heritability (broad sense) estimates were high (>70%) for spike length, biological yield per spike and number of grains per spike among all the three generations. In BC_1_F_3_ and F_4_ generations highest heritability was recorded for spike length while in BC_2_F_2_, number of grains per spike showed highest heritability. Genetic advance as 5% of mean indicated a good scope for improvement for biological yield per plant and harvest index in all the generations. However, in BC_1_F_3_, number of tillers per plant and in BC_2_F_2_ and F_4_ grain yield per plant recorded highest value of genetic advance as 5% of mean.

**Table 3:**
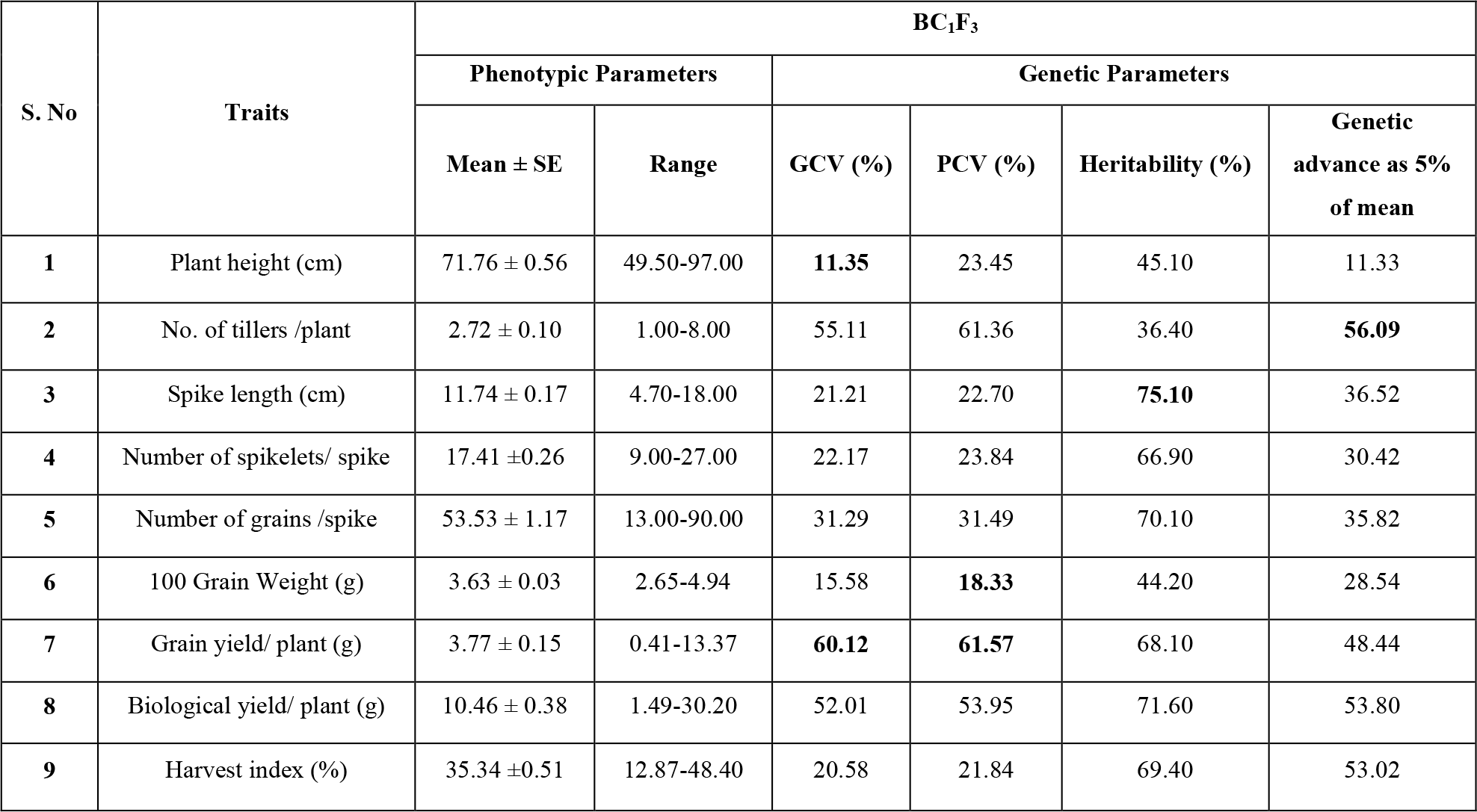
Mean, range, phenotypic and genotypic coefficient of variation, heritability and genetic advance for various traits in BC_1_F_3_ population.

**Table 4:** Mean, range, phenotypic & genotypic coefficient of variation, heritability and genetic advance for various traits in BC_2_F_2_ population.

| S. No | Traits | BC<sub>2</sub>F<sub>2</sub> |  |  |  |  |  |
| --- | --- | --- | --- | --- | --- | --- | --- |
|  |  | Phenotypic Parameters |  | Genetic Parameters |  |  |  |
|  |  | Mean ± S.E | Range | GCV (%) | PCV (%) | Heritability (%) | Genetic advance as 5% of mean |
| 1 | Plant height (cm) | 73.17±1.05 | 53.50-89.50 | **15.15** | 40.6 | 23.9 | 11.64 |
| 2 | No. of tillers /plant | 2.27±0.16 | 1.00-8.00 | 44.72 | 55.86 | 37.7 | 56.54 |
| 3 | Spike length (cm) | 10.22±0.26 | 4.50-15.70 | 18.93 | **21.39** | 76.3 | 34.51 |
| 4 | Number of spikelets/spike | 14.68±0.488 | 5.00-23.00 | 16.51 | 18.74 | **77.6** | 29.94 |
| 5 | Number of grains/spike | 52.63±1.88 | 15.00-84.00 | 35.52 | 35.65 | 75.2 | **62.84** |
| 6 | 100 Grain Weight (g) | 3.49±0.09 | 0.90-4.76 | 21.2 | 31.78 | 44.5 | 29.13 |
| 7 | Grain yield/ plant (g) | 3.89±0.37 | 0.36-14.17 | **51.79** | **55.94** | 68.3 | 55.24 |
| 8 | Biological yield/ plant (g) | 10.03±0.79 | 2.20-31.50 | 42.98 | 44.86 | 71.8 | 54.82 |
| 9 | Harvest index (%) | 37.00±0.95 | 16.36-46.59 | 27.43 | 28.06 | 75.6 | 55.25 |

**Table 5:** Mean, range, phenotypic & genotypic coefficient of variation, heritability and genetic advance for various traits in F_4_ population.

| S. No | Traits | F<sub>4</sub> |  |  |  |  |  |
| --- | --- | --- | --- | --- | --- | --- | --- |
|  |  | Phenotypic Parameters |  | Genetic Parameters |  |  |  |
| | | Mean ± SE | Range | GCV (%) | PCV (%) | Heritability (%) | Genetic advance as 5% of mean |
| 1 | Plant height (cm) | 78.33±1.36 | 42.60-105.50 | **14.77** | 21.14 | 45.10 | 13.93 |
| 2 | No. of tillers /plant | 3.65±0.19 | 1.00-8.00 | 43.98 | 53.04 | 27.82 | 56.09 |
| 3 | Spike length (cm) | 10.12±0.23 | 6.10-19.60 | 19.81 | 22.13 | **75.10** | 36.52 |
| 4 | Number of spikelets/ spike | 14.51±0.28 | 7.00-19.00 | 16.84 | 19.21 | 66.90 | 30.42 |
| 5 | Number of grains /spike | 45.50±1.74 | 4.00-80.00 | 32.53 | 32.69 | 70.10 | 29.82 |
| 6 | 100 Grain Weight (g) | 3.63±0.07 | 2.84-4.97 | 14.92 | 18.31 | 44.20 | 28.54 |
| 7 | Grain yield/ plant (g) | 3.81±0.22 | 0.62-8.60 | **49.12** | **54.24** | 68.10 | **58.44** |
| 8 | Biological yield/ plant (g) | 10.47±0.52 | 2.90-20.90 | 21.23 | 43.42 | 71.60 | 53.80 |
| 9 | Harvest index (%) | 36.12±1.01 | 13.26-48.60 | 23.35 | 33.97 | 69.40 | 53.02 |

### Screening of backcrosses and F_4_ populations using markers for Nax loci

Primers for *Nax1* and *Nax2* were used to amplify genomic DNA of parents. These primers gave amplification only in Kharchia 65. A total of 150 plants (Table 2) of all the population were screened for both *Nax* loci. Thirty-five plants confirmed the presence of *Nax1* locus at a band size of 210 bp while only 24 plants confirmed the presence of *Nax2* locus at a band size of 225 bp. Fifteen plants were found to have both the *Nax* loci.

### Background selection in selected backcrosses and F_4_ plants

A total of 178 SSR primers were used to study the polymorphism, out of which 30 SSR primers showed polymorphism between the two parents (Table 6). These polymorphic primers were then used for background selection in 44 phenotypically superior and selected plants (BC_1_F_3_ - 24, BC_2_F_2_-1 and F_4_ −19). Out of these 44 plants, 15 were having both *Nax1* and *Nax2* loci, 20 had only *Nax1* locus and 9 had only *Nax2* locus. Polymorphism information content (PIC) value gives the information about the amount of variability for a primer in the studied plants and it ranged from 0.035 (*wmc*145) to 0.647 (*Xgwm*356) with an average value of 0.350 for all generations of cross. The PIC value of primers for A-genome ranged 0.035-0.647 with an average of 0.338. For B and D genome, PIC value ranged 0.057 - 0.482 and 0.301 - 0.491 with an average of 0.318 and 0.380, respectively.

**Table 6:**
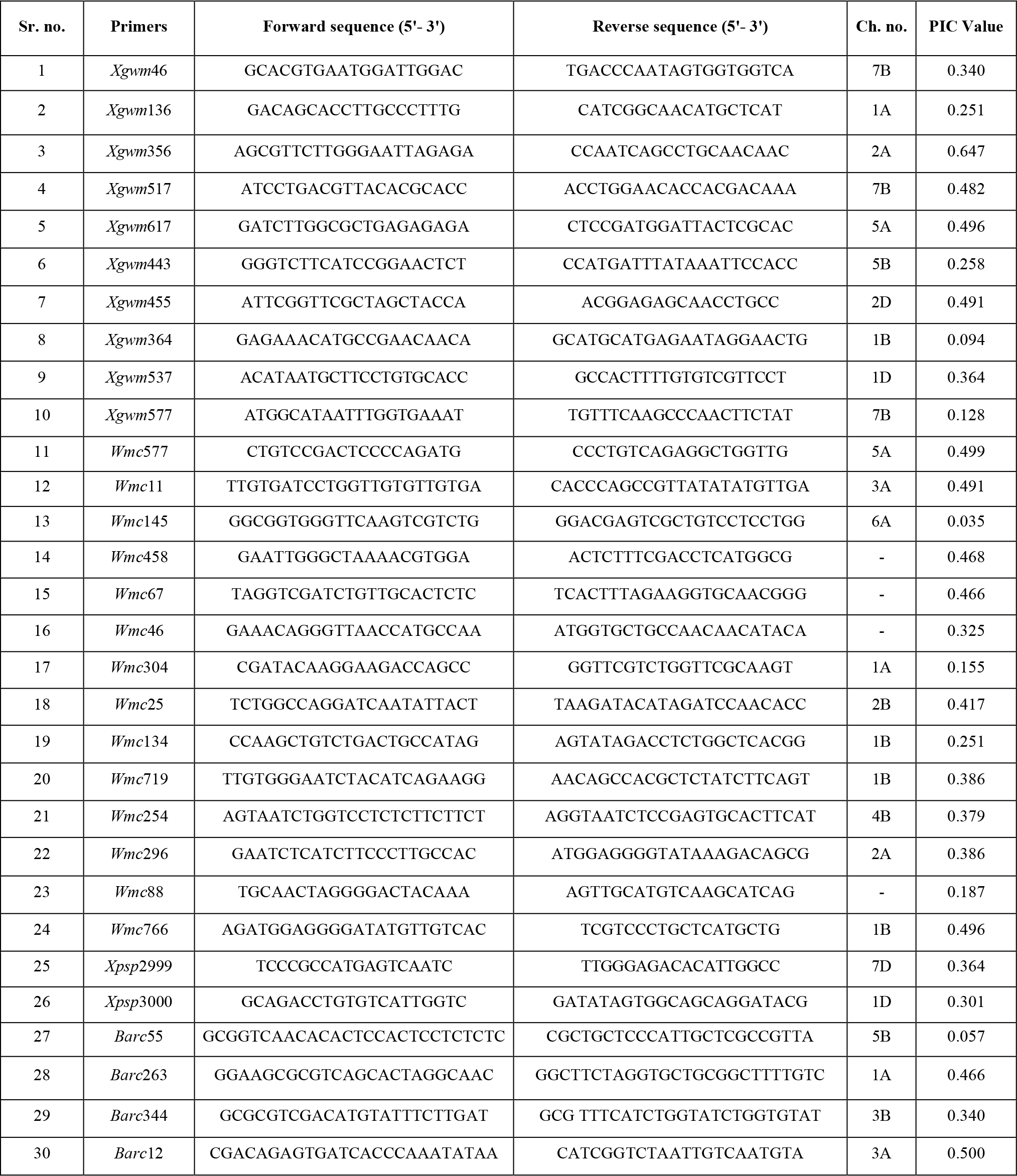
A brief description of polymorphic SSRs.

### Genetic relationship and cluster tree analysis

Two major clusters were formed at a similarity coefficient of 0.42 and 0.45 for BC_1_F_3_ and F_4_ respectively. Since in BC_2_F_2_ only one plant showed positive result for *Nax1* locus so this generation was not considered for background selection and further evaluation. For evaluated populations, major cluster 1 was having WH 1105 along with the progeny plants and the other, major cluster II was having only Kharchia 65 (Fig. 1 and 2). This clustering was also evident from the two dimensional principle component analysis (PCA) (Fig. 3 and 4). The result is coherent with the dendrogram generated employing UPGMA and is a further confirmation of genetic similarities delineated in the present study.

**Figure 1:**
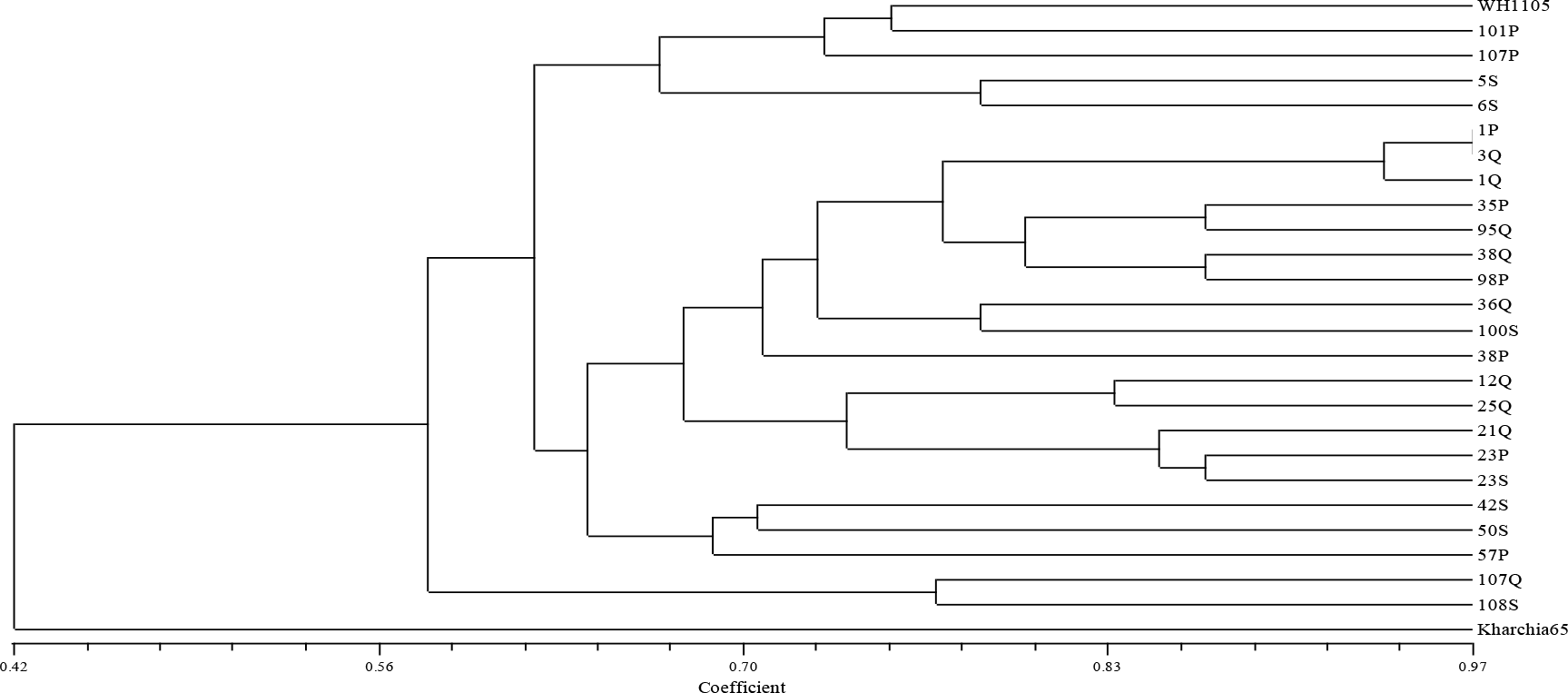
Dendrogram showing relationship among parents and BC_1_F_3_ plants at 30 polymorphic SSR loci.

**Figure 2:**
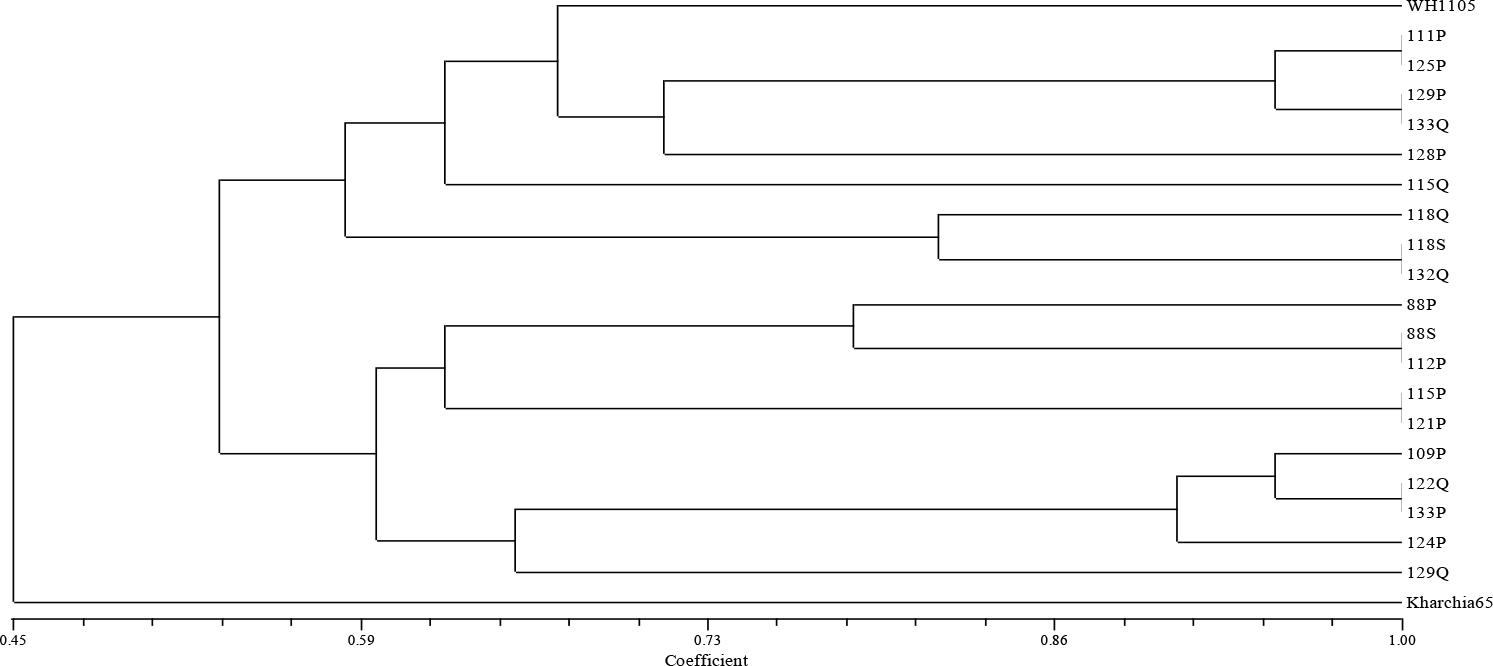
Dendrogram showing relationship among parents and F_4_ plants at 30 polymorphic SSR loci.

**Figure 3:**
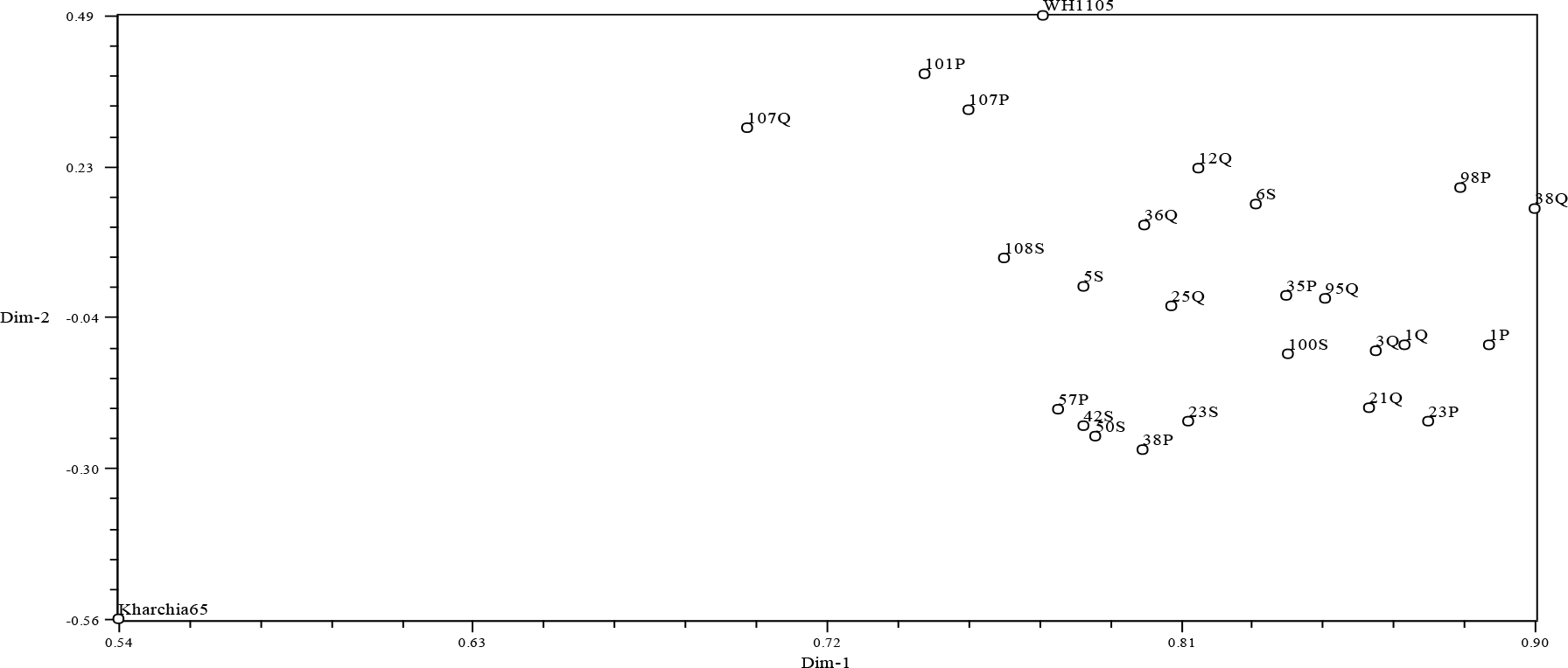
Two dimensional PCA scaling of parents and BC_1_F_3_ plants.

**Figure 4:**
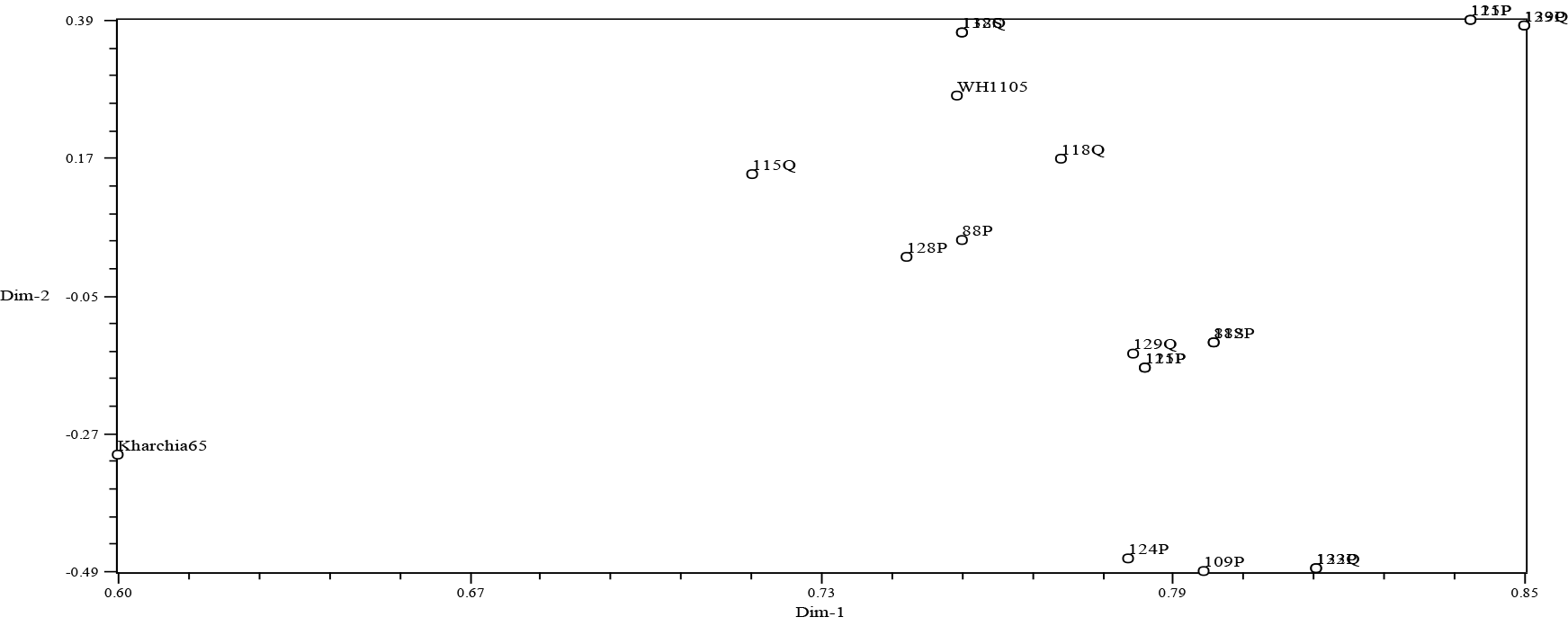
Two dimensional PCA scaling of parents and F_4_ plants.

## DISCUSSION

Wheat (*Triticum aestivum* L.) is one of most important crop plants worldwide which suffers significant grain yield losses due to soil salinity. Salinity variously affects the plants like, germination vigor, plant height, leaf area, number of effective tillers per plant, number of spikes per plant, dry shoot weight and grain yield (Dadrwal, 2018). Although, there are several strategies to increase wheat production in the salt-affected areas (such as leaching and drainage), the cultivation of tolerant genotypes is recognized as the most effective way to overcome this limitation. The cultivar, Kharchia 65, is one of the very few reputed donors of salt tolerance in wheat and has been extensively used in breeding for salt tolerance cultivars globally. However, attempts to improve the salt tolerance through conventional breeding programmes have met with very limited success due to complexity of the trait which is governed by several physiological and genetic factors, and is growth stage specific (Haq *et al.*, 2010). Marker assisted selection is a promising technique that brings genes of interest into plants. Two genes, *Nax1 and Nax2* for Na^+^ exclusion originating from *T. monococcum* have been validated to be present in Kharchia 65 (Yadav *et al.*, 2017). However, in practice relying completely on linked markers is not enough, as there is not much information about all of the available alleles of all the genes, and their interactions with each other (dominance for alleles at a single locus, and epistasis for alleles at different loci), with genetic background, and the environment. Germination indices along with some agronomic traits and their relationship with salt tolerance indices could be a feasible means for selection of salt tolerance (Diaguna, 2017; Hasan *et al.*, 2017).

### Evaluation of parents, backcrosses and F_4_ populations under salt and normal condition

The comparative evaluation of parents revealed significant reduction in the various trait values for all the traits in WH 1105 as compared to Kharchia 65 under saline conditions. Turki *et al.* (2012) also revealed significant reduced values for various traits in their results while investigating effect of salinity on grain yield and quality in 55 genotypes of durum and common wheat. A comparison of the backcrosses and F_4_ generations with the parents under initial salinity stress depicted that BC_1_F_3_ and BC_2_F_2_ generation plants showed more improvement as compared to F_4_ plants, showing that backcrossing has led to accumulation of genes for high yield and the selection method used so far is effective. High GCV and PCV were observed for grain yield per plant under salinity conditions. Similar findings were reported earlier by Naik *et al.* (2015) and Kumar *et al.* (2013, 2014) while studying the genetic variability parameters and heritability in bread wheat. The PCV was considerably higher than the GCV for plant height and biological yield in F_4_, for plant height in BC_1_F_3_ and for plant height, number of tillers per plant and 100 grain weight in BC_2_F_2_ generation. Similarly, Ali *et al.* (2008) reported considerable higher PCV than GCV for yield per plant (51.27 and 41.19, respectively) and number of productive tillers per plant (35.24 and 23.79, respectively). This shows that these traits were more affected by environmental effects than other traits. Comparatively high heritability was recorded for spike length and number of grains per spike in all the three generations. The results are in agreement with previous studies of Ijaz and Shahzad (2015) evaluated important quantitative traits for F_2_ progenies and found highest heritability for spike length and number of spikelets per spike. Ahmed *et al.* (2016) reported higher heritability estimates for plant height, number of tillers per plant, 1000- grain weight and grain yield per plant for the entire cross combinations and Safi *et al.* (2017) found higher heritability across the years for plant height, 100 grain weight and grain yield. Traits with high heritability values can be improved directly through selection since these traits are less influenced by environment and there would be greater correspondence between phenotypic and breeding values (Panse, 1957).

In all the generations, high genetic advance as 5 % of mean was observed for number of tillers per plant, grain yield per plant, biological yield per plant and harvest index. Kumar *et al.* (2013) also found higher genetic advance for plant height, biological yield per plant and harvest index. Ahmed *et al.* (2016) estimated higher genetic advance for plant height, number of tillers per plant, spike length, number of grains per spike, 1000-grain weight, and grain yield per plant in his study of estimating the inheritance of yield components in wheat gown in drought condition. High heritability and genetic advance are attributed to additive gene action which would result in effective selection for these traits. High heritability and low genetic advance was reported for plant height across all the generations under study. High heritability and low genetic advance is attributed to non-additive gene action. For such traits, hybridization would result in effective utilization of variation. Rangare *et al.* (2010) recorded high heritability and low genetic advance for plant height While, Kumar *et al.* (2014) and Ahmad et al. (2016) found high heritability and high genetic advance for plant height in seven F_3_ progenies and their 8 parental lines. A comparison of BC_1_F_3_, BC_2_F_2_ and F_4_ plants with parents under initial salt stress conditions revealed that there was improvement in all the traits.

### Molecular marker analysis

The use of genetic and genomic analysis to identify DNA regions tightly linked to quantitative traits in crops, called “molecular marker-assisted breeding”, can facilitate breeding strategies for wheat improvement (Munns and Tester, 2008). Marker assisted selection approach is more effective because selection based on phenotypic traits is restricted by variation in soil salinity and sodicity (Ma *et al.*, 2007). Microsatellites or SSRs as powerful genetic markers have been extensively used for investigation of genetic divergence, genome mapping and QTL mapping for salt tolerance in different crops (Liu *et al.*, 2001; Munns *et al.*, 2002; Ma *et al.*, 2007 and Kurup *et al.*, 2009).

In the present study, DNA isolated from 150 phenotypically superior plants was analyzed for *Nax1* and *Nax2* loci. Out of these 150 plants, 35 plants (18 BC_1_F_3_plants, 1 BC_2_F_2_ plant and 16 F_4_ plants) confirmed the presence of *Nax1* locus at 210 bp band size and 13 BC_1_F_3_ plants and 11 F_4_ plants showed presence of *Nax2* locus at 225 bp band size. 15 plants comprising of 7 BC_1_F_3_ and 8 F_4_ plants of WH 1105 x Kharchia 65, were found to have both the *Nax* loci. A total of 43 plants [BC_1_F_3_ (24) and F_4_ (19)] were selected for further molecular analysis. Among selected 44 plants, 15 were having both the *Nax* loci, 20 had only *Nax1* locus and 9 had only *Nax2* locus. Agarose gel electrophoresis confirmed *Nax1* and *Nax2* loci with presence of band size of 210 bp and 225 bp respectively and similar banding pattern of *Nax* loci was found in Turkey wheat accessions (Abbasov *et al.* 2011). De Bustos *et al.* (2001) also developed and used gene specific markers to assist selection in backcross progenies to improve the glutenin quality in spanish wheat. The selection was also assisted using other polymorphic systems (AFLPs) in recovering the genetic background of the recurrent parent. Vishwakarma *et al.* (2014) obtained similar results while introgressing a high grain protein gene *Gpc-B1* in an elite wheat variety of Indo-Gangetic Plains through marker assisted backcross breeding. James *et al.* (2012) had similar results in bread wheat after introgression of salinity tolerance genes from durum wheat.

30 polymorphic SSR markers provided uniform coverage of all three wheat genomes (10, 12 and 4 markers on A, B and D chromosome, respectively) in selected 44 salt tolerant plants of WH 1105 x Kharchia 65. Ten polymorphic markers were located on D genome since Gorham et al. (1987) reported that salinity tolerance is controlled by D genome. Genc *et al.* (2010) reported five QTLs present in bread wheat for Na^+^ exclusion and observed improvement in performance under salinity. The NTSYS-pc UPGMA tree cluster analysis and two dimensional PCA scaling revealed that salt tolerant wheat variety Kharchia 65 was quiet distinct and diverse from WH 1105 based on their similarity matrix. All the plants were inclined towards recurrent parent WH 1105 representing higher similarity with the recurrent parent. Kharchia 65 was clustered separately from the recurrent parent and progenies, indicating high dissimilarity between them.

### Selection of promising plants

On the basis of presence of *Nax* loci, background selection and performance in initial salt stress, selection of plants from all the generations was made and selected plants were further taken to the next generation trial which would stabilize genome of recurrent parent with salt tolerant genes from the donor parent in successive lines/ progenies.

## CONCLUSION

On the basis of evaluation of allelic profile at *Nax1* and *Nax2* loci, polymorphic SSR markers, net house evaluation under initial salt stress condition and higher grain yield (comparable to respective salinity sensitive wheat parent) led to the identification of 44 plants of cross. Four plants, 2 (BC_1_F_3_), 1 (BC_2_F_2_) and 1 (F_4_) were selected due to their better mean performance and high heritability coupled with high genetic advance in most of the characters under salinity condition. These plants could be further backcrossed with the recurrent parent WH 1105 to develop salt tolerant wheat lines. The best breeding strategy is to select for high yield on non-saline soils and it certainly follows that in a backcrossing breeding programme, the most productive genotype at the lowest salinity should always be used as the recurrent parent. However, plants were selected under salt stress in our study. It clearly indicates that a linked marker like *Nax1* and *Nax2* could provide a valuable tool for breeding wheat with enhanced tolerance to salinity conditions.

